# Spatial transcriptomics reveals influence of microenvironment on intrinsic fates in melanoma therapy resistance

**DOI:** 10.1101/2024.06.30.601416

**Authors:** Ryan H. Boe, Catherine G. Triandafillou, Rossana Lazcano, Jennifer A. Wargo, Arjun Raj

## Abstract

Resistance to cancer therapy is driven by both cell-intrinsic and microenvironmental factors. Previous work has revealed that multiple resistant cell fates emerge in melanoma following treatment with targeted therapy and that, *in vitro*, these resistant fates are determined by the transcriptional state of individual cells prior to exposure to treatment. What remains unclear is whether these resistant fates are shared across different genetic backgrounds and how, if at all, these resistant fates interact with the tumor microenvironment. Through spatial transcriptomics and single-cell RNA sequencing, we uncovered distinct resistance programs in melanoma cells shaped by both intrinsic cellular states and the tumor microenvironment. Consensus non-negative matrix factorization revealed shared intrinsic resistance programs across different cell lines, highlighting the presence of universal and unique resistance pathways. In patient samples, we demonstrated that these resistance programs coexist within individual tumors and associate with diverse immune signatures, suggesting that the tumor microenvironment and distribution of resistant fates are closely connected. Single-cell resolution spatial transcriptomics in xenograft models revealed both intrinsically determined and extrinsically influenced resistant fates. Overall, this work demonstrates that each therapy resistant fate coexists with a distinct immune microenvironment in tumors and that, *in vivo*, tissue features, such as regions of necrosis, can influence which resistant fate is adopted.

## Introduction

Many aspects of cancer biology, such as oncogenesis, therapy resistance, and metastasis are driven by the behavior of individual cells^1–9^. Often, these outcomes result from mutations in rare cells^10^; however, it has become increasingly clear that non-genetic differences can also be responsible for these phenomena^11^. Recent evidence has shown that these non-genetic differences can come in a spectrum, potentially leading to multiple outcomes as cells transition to becoming, for instance, therapy-resistant^12–16^. The implication is that cell-intrinsic differences can be amplified during the acquisition of therapy resistance. However, it is also clear that *in vivo* interactions with the local microenvironment, including the immune system, can also shape the outcome of therapy resistance, metastasis, and more. The relative contributions of cell-intrinsic and cell-extrinsic factors to how cells become resistant *in vivo* remains largely unknown.

Studies of cell-intrinsic factors contributing to therapy resistance have demonstrated that “twin” cells that share the same non-genetic state will typically adopt the same fate when separately subjected to the same external stimulus and that different non-genetic initial states can lead to different outcomes^13,17–19^. Tracking individual clones longitudinally using DNA barcoding has been essential to reaching these conclusions about the shared fates of “twin” cells and the importance of initial non-genetic states to ultimate fates. In Goyal et al., these techniques showed that individual resistant colonies, formed *in vitro* in homogeneous conditions, display remarkable differences in their morphological, molecular, proliferative, and invasive properties, leading to the conclusion that cell-intrinsic differences can lead to the formation of different resistant “fates”^13^. However, these studies were primarily done in cell culture in highly controlled conditions. It remains to be seen whether these resistant fates appear in the more complex microenvironments that occur *in vivo*.

At the same time, it is also clear that, *in vivo*, there is extrinsic instruction of cell fates via interactions with stromal cells^20,21^, extracellular matrix^22^, and cells from the immune system^23,24^. We currently do not know how these extrinsic cues influence intrinsically specified resistant fates. It is possible, for instance, that different resistant fates can only emerge in different areas of the resistant tumor or that only certain immune cell types are able to infiltrate or otherwise interact with certain resistant fates. Testing for these possibilities requires the use of spatial analysis methods, along with new analytical methods to discriminate between both intrinsic and extrinsic cell fate specification.

Here, we used spatial transcriptomics datasets to look for signatures of diverse resistant fates in patient samples. We integrated multiple *in vitro* single-cell datasets to arrive at consensus markers of resistant fates and then looked for those fates in therapy-resistant melanoma, finding that these fates exist in patients and that they are correlated with various immune signatures (Figure 1A). We also analyzed xenograft models of therapy-resistant melanoma with single-cell resolution, finding resistant fates that were intrinsically determined, fates that were extrinsically determined, and fates that had some component of extrinsic and intrinsic determination. Overall, we found that intrinsic melanoma resistance fates are associated with the local microenvironment, suggesting that both intrinsic and extrinsic factors need to be considered when predicting resistant fate specification.

**Figure 1.**
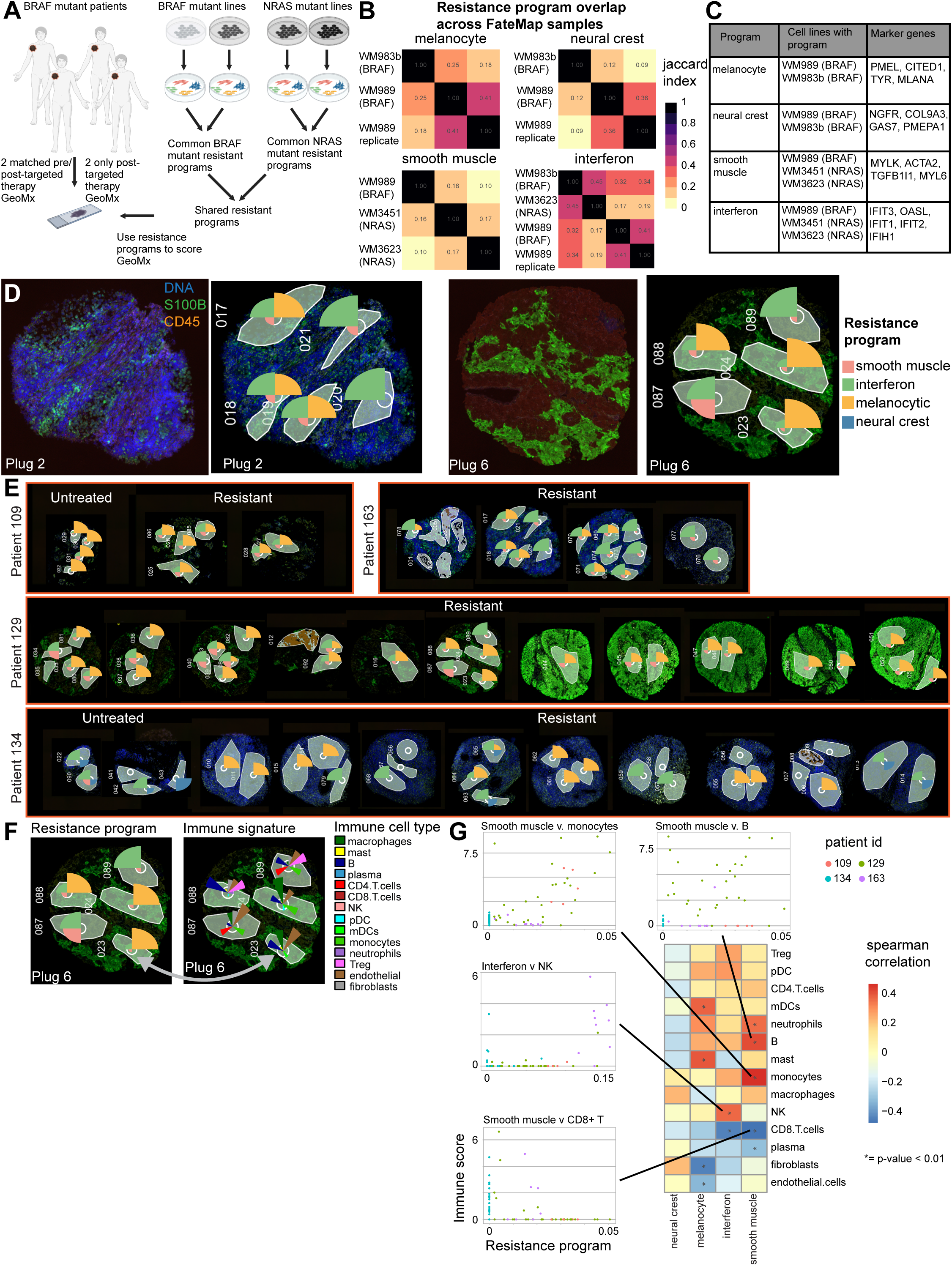
GeoMx spatial transcriptomics reveals distribution of intrinsic melanoma resistance programs and their immune associations in patient samples. A. Schematic of experimental set-up and computational strategy to identify shared resistance programs. We spatially profiled using the GeoMx system samples from four patients total, two with matched pre-/post-targeted therapy and one with only post-targeted therapy. Separately, we barcoded, treated with targeted therapy, and single-cell sequenced four melanoma cell lines, two with *BRAF* mutations and two with *NRAS* mutations. From each separate single-cell dataset, we found resistance programs using cNMF and then compared them to find shared resistance programs across cell lines and mutations. B. Heatmaps showing the Jaccard index for each intrinsic resistance program compared across cell lines. We identified recurrent gene programs using cNMF. We then examined the top genes for each NMF program and assigned a function to that program. We then calculated the jaccard index for the top 100 marker genes for each pairwise combination of resistance programs across all cell line samples (see supplementary figure 1 for the full heatmap). We then identified metaprograms as programs that were recurrent across multiple samples with a jaccard index of at least 0.10. C. Table with each intrinsic resistance program, the cell lines that displayed that program, and the top consistent marker genes for the program. D. Two sets of patient tumor plugs with staining alone (left images) and with region of interest (ROI) and resistance program score drawn (right images). Samples are stained for DNA (SYTO 13, blue), S100B (green), and CD45 (red). For each ROI, the total gene expression profile was deconvolved using a signature matrix of the consensus metaprograms using non-negative least squares. The scores for the four intrinsic resistance programs were normalized to one for each ROI, and the area of each wedge is proportional to its relative resistance program score. E. Intrinsic resistance program scores for each ROI across all four patients. Intrinsic resistance program scoring is the same as in panel D. F. Comparison between the melanoma resistance program score and the immune cell signature for each tumor-containing ROI. For each tumor-containing ROI, we estimated the immune cell types present using the SpatialDecon package from NanoString (right image). We can then compare, within each ROI, which resistance programs are expressed (left image) to the immune cell infiltration in the ROI. G. Heatmap with correlations between resistance program score and immune cell score in resistant sample, tumor-containing ROIs (bottom right) and individual scatter plots for resistance program scores versus immune signature scores (top and left). We calculated the Spearman correlation between the melanoma program score and immune cell score across all post-treatment, tumor ROIs and marked those with p-values < 0.01. We pulled scatter plots for four associations that were statistically significant where each dot is a single ROI and is colored by which patient the ROI originated from.

## Results

### Consensus Non-negative Matrix Factorization of FateMap samples reveals shared intrinsic resistance programs

Before analyzing patient samples for the presence of resistant fates, we wanted to systematically define a gene program for each resistant fate. Since resistant fates were defined from *in vitro* data, our approach was to develop a consensus resistant-fate signature using all of our available *in vitro* datasets. In our previous work^13^, we used FateMap to reveal that a number of intrinsically determined resistant fates arise in clonal *BRAF* mutant melanoma cells upon exposure to MAPK inhibition. In Goyal et al., we uniquely barcoded melanoma cells, allowed them to divide, and then split cells into multiple arms on the basis that sister cells with the same barcode retain similar transcriptomic profiles and phenotypic properties^12,13,25^. We then treated both arms with *BRAF*^V600E^ inhibitor to create resistant cells and found the unique resistant fate each cell adopted. We then checked whether sister cells adopted the same resistant fate, which would indicate that the state of the cell before being treated with drug determined the resistant fate, i.e. the resistant fate was intrinsically determined. In FateMap, we found five resistant fates–*MLANA* high, *NGFR* high, *ACTA2* high, *IFIT2* high, and *VCAM1* high–that were intrinsically determined. A limited analysis suggested that some number of marker genes for each resistant fate are shared with two *BRAF*^V600E^ lines (WM989 and WM983b) and even in two *NRAS*^Q61K^ lines (WM3251 and WM3623). However, we did not fully assess the similarity between the resistant fates seen in each FateMap sample. We reasoned that resistant fates that recur between multiple cell lines were more likely to reflect general resistant fates that might appear in patient data.

To determine the level of overlap of resistant fates across cell lines, we used consensus non-negative matrix factorization (cNMF) to first find gene expression programs independently for each FateMap sample. cNMF attempts to decompose the single-cell cell-by-gene expression matrix into two new matrices, a program-by-gene signature matrix and a cell-by-program expression matrix^26^. The first matrix gives a number of programs for which each gene has some expression value. For each program, the genes with the top expression, or the top unique expression compared to other programs, can be used to infer the identity of the program. The second matrix gives the amount of each program that each cell expresses. A major advantage of this method compared to traditional clustering methods is that each cell can express multiple programs and this expression is quantitative rather than the single, discrete cluster assignment given by traditional methods. As opposed to clustering, which is often treated as the cell’s “identity,” some programs may capture a cell’s identity, whereas others capture what activity the cell was performing at the time of sequencing. Another advantage of our approach is that by finding gene expression programs independently for each sample and then comparing the programs, we minimize the impact of technical variation and avoid the fraught problem of dataset integration. We checked for recurrent programs across samples by calculating the Jaccard index, which calculates the amount of overlap between two sets of genes, for the top markers for each program and performing hierarchical clustering (Supplementary Figure 1). Our ultimate goal was to identify programs that were represented among multiple FateMap experiments, so we defined a “metaprogram” as a set of cNMF programs that overlap with a Jaccard index of at least 0.1 in at least two FateMap samples. This analysis yielded 13 expression metaprograms that recurred in at least two FateMap samples. Reassuringly, we found that a number of the metaprograms corresponded to well-defined cellular activities that have been previously reported in pan-cancer NMF analyses^27–29^, such as cell division and epithelial-to-mesenchymal transition. In addition, one metaprogram corresponded to high mitochondrial gene expression, which we took to signify damaged and dying cells.

We found that while many metaprograms were shared between cell lines, others were specific to certain cell lines or to lines that shared a particular driver mutation. From these 13 metaprograms, we recapitulated four of the resistant fates previously identified in FateMap–the melanocyte (*MLANA* high), neural crest (*NGFR* high), smooth muscle (*ACTA2* high), and interferon (*IFIT2* high) metaprograms (Figure 1B). The remaining nine metaprograms were not identified as intrinsically-determined resistance programs in FateMap, which we took to mean they were either cell-activity programs or resistant fates that were not identified in the previous FateMap analysis. The *VCAM1* high fate from FateMap did not appear as a cNMF program in any sample, suggesting that the program may be too weak or too rare to find by this method. The melanocyte and neural crest resistance programs did not occur in the *NRAS* mutant FateMap samples, potentially suggesting that these two resistant fates are specific to *BRAF*^V600E^ mutant melanoma. Alternatively, the lack of the melanocyte and neural crest resistance programs in the *NRAS*^Q61K^ lines may reflect the fact that the *NRAS*^Q61K^ mutant samples were treated with the MEK inhibitor trametinib while the *BRAF*^V600E^ samples were treated with *BRAF*^V600E^ inhibitor vemurafenib, which we have shown decreases the proportion of the *MLANA* high resistant fate^13^. We also found that the smooth muscle and interferon programs were absent from WM983b *BRAF*^V600E^ mutant line. Next, we took the intersection of the marker genes for each resistance program to identify a consensus marker gene set for each. These are the key marker genes that we used to look for metaprogram expression in patient samples. While many of the genes identified in this manner were the same as those previously identified by single-cell clustering, our method identified unique top markers that were shared across samples, such as *PMEPA1* for the neural crest program, *TGFB1I1* and *MYL6* for the smooth muscle program, *CITED1* and *TYR* for the melanocyte program, and *IFIT3*, *IFIT1*, and *IFIH1* for the interferon program (Figure 1C). Together, these results show that very different cell lines share some programs, while others are unique to each cell line and that there is a consensus set of marker genes for each resistance program across cell lines. These results further suggest that genetic background and driver mutation may influence which resistant fates emerge upon treatment.

### Spatial transcriptomics reveals intrinsic resistance programs in patient samples

Having identified a consensus gene set for each metaprogram, we next asked whether there was any evidence for these metaprograms in patient samples using our recently published GeoMx dataset (Supplementary Figure 2). Our previous analysis revealed that several key resistance markers were differentially expressed in different regions of the same tumor; however, we did not quantify expression beyond the level of individual marker genes. We used non-negative least squares deconvolution to quantify the amount of gene metaprogram expression in each region (Figure 1D). We performed the deconvolution with 12 of the 13 metaprograms (we removed the mitochondrial metaprogram, which represented poor-quality cells) (Supplementary Figure 3). Importantly, since some genes expressed by resistant melanoma cells, including key markers of resistant fates, are also expressed tumor stromal and immune cells, we excluded metaprogram genes that were known to be expressed in cells of the tumor microenvironment (see methods for details). In addition to deconvolving the resistant samples, we also performed the deconvolution on the pre-treatment samples, even though the metaprograms were derived from resistant cell expression. We determined the goodness of fit of the deconvolution by calculating the log-likelihood of the deconvolution assuming a negative binomial model for the expression of each gene in the deconvolution, where the mean for each gene was the predicted value calculated by multiplying the signature matrix by the transpose of the deconvolution matrix, essentially reversing the deconvolution and assessing how close this predicted value was to the true expression value. We removed instances where the predicted value was zero since a mean of zero is not possible with a negative binomial model. Since we had no estimate of the variance, we used three different values and compared the results. We then calculated a null distribution using a permutation test. We shuffled the row labels (genes) of the signature matri× 1000 times, deconvolved the data, and then performed the log-likelihood calculation described above with the three variance values. We then calculated the total log-likelihood for each region and compared it to the total log-likelihood for each region under the null distribution. We found that our deconvolution was better than shuffled null for 92 out of 95 regions at a p-value cutoff of 0.01 (Supplementary Table 1). Overall, we were satisfied that the deconvolution was of sufficient quality to look at the distribution of resistance metaprograms in our patient samples.

We decided to focus on the results from the four resistance metaprograms that corresponded to intrinsically determined fates in the original FateMap analysis, the melanocyte, neural crest, smooth muscle, and interferon metaprograms. We found that all but seven of the 92 regions had non-zero resistance program expression, confirming the presence of resistance programs in patient samples (Figure 1E). We found that while there was a patient-specific signature to resistance program expression, there were also many examples of adjacent regions from the same biopsy plug with very different resistance program expression, suggesting that multiple resistant fates can emerge within patients, albeit with patient-specific biases in the particular resistant fates that do emerge (Figure 1E). This patient-specific bias can even be seen in pre-treatment samples, with patient 109 showing high melanocyte program scoring pre-treatment while patient 134 showed no melanocyte scoring pre-treatment. The scoring of resistance programs in pre-treatment samples likely reflects some degree of cell state heterogeneity in the drug-naive population, as has been shown by others^30^, and that these drug-naive transcriptional programs partially overlap with our resistance programs. However, the fact that different regions of the same tumor display different resistance programs suggests some degree of intrinsic fate determination. It is important to note that these resistant cell fate differences observed in tissue may reflect genetic heterogeneity throughout the tumor; our data show that it is consistent with the non-genetic variability observed in the clonal cell lines originally observed in FateMap. Overall, our analysis suggested that multiple resistant fates emerge within a single patient, although each patient may have a bias towards a particular distribution of these fates.

### Intrinsic resistance programs are associated with the tumor immune microenvironment

We were curious whether we could find evidence of specific interactions between particular resistance programs and the tumor microenvironment. We performed an independent deconvolution analysis on the same GeoMx dataset using a tumor immune signature (Supplementary Figure 4). We therefore were able to compare, for each region, the resistance program score and the immune signature score and look for associations between them (Figure 1F). We limited our analysis to resistant sample regions that were annotated as containing predominantly tumor cells by a pathologist (Supplementary Table 2). We found multiple associations between the melanocytic, smooth muscle, and interferon resistance programs and the tumor microenvironment (Figure 1G). In particular, there was a positive association between interferon program expression and natural killer (NK) cell score, though the association was mostly driven by a single patient. We also found that the smooth muscle program was positively associated with B cell infiltration but negatively associated with CD8+ T cell infiltration, suggesting that resistant fate may influence or be influenced by the makeup of tumor-infiltrating lymphocytes in a region. The smooth muscle program had, of all the resistant fates, the most positive associations with the enrichment of different immune cell types, suggesting that it may be the most immunogenic. No associations reached statistical significance for the neural crest program, likely due to the small number of regions that contained this program. A statistically significant, positive association with macrophage infiltration emerged if we included pre-treatment regions in the analysis (Supplementary Figure 5).

### Single-cell spatial transcriptomics reveals intrinsically and extrinsically determined resistant fates

We next wanted to look for spatial patterns of expression in targeted therapy-resistant melanoma at the single-cell level. To do so in a more precisely controlled environment than human samples, we used the Spatial Genomics platform with a custom panel of 427 genes (Supplementary Table 3) to spatially profile single cells of clonal *BRAF* mutant WM989 A6-G3 5a3 cells that were implanted into a mouse xenograft and then made resistant via treatment with targeted therapy. Unsupervised clustering of single cells using the leiden algorithm, a community detection algorithm that iteratively assigns cells into “communities” based on connectivity in PCA space, revealed seven clusters (Figure 2A). These seven included clusters that clearly mapped to the melanocyte and the interferon resistance program found from *in vitro* FateMap (Figure 2B). There additionally was a cluster that expressed *COL9A3* and *S100B* and was weakly enriched for *NGFR*, though these genes were less confined to a single cluster than *in vitro* (Supplementary figure 6). Notably, we did not find the smooth muscle state, likely partially because the *ACTA2* probe, a key marker of the state, failed to hybridize for technical reasons. Uniquely, compared to *in vitro*, we also found a cluster highly enriched for *JUNB and FOS*. This cluster likely corresponds to the “stress-like” state previously described in zebrafish models of melanoma, which also showed high expression of *JUNB* and *FOS*^31^ (Figure 2B). Finally, we found a number of clusters that did not clearly map to a resistance fate (Figure 2B).

**Figure 2.**
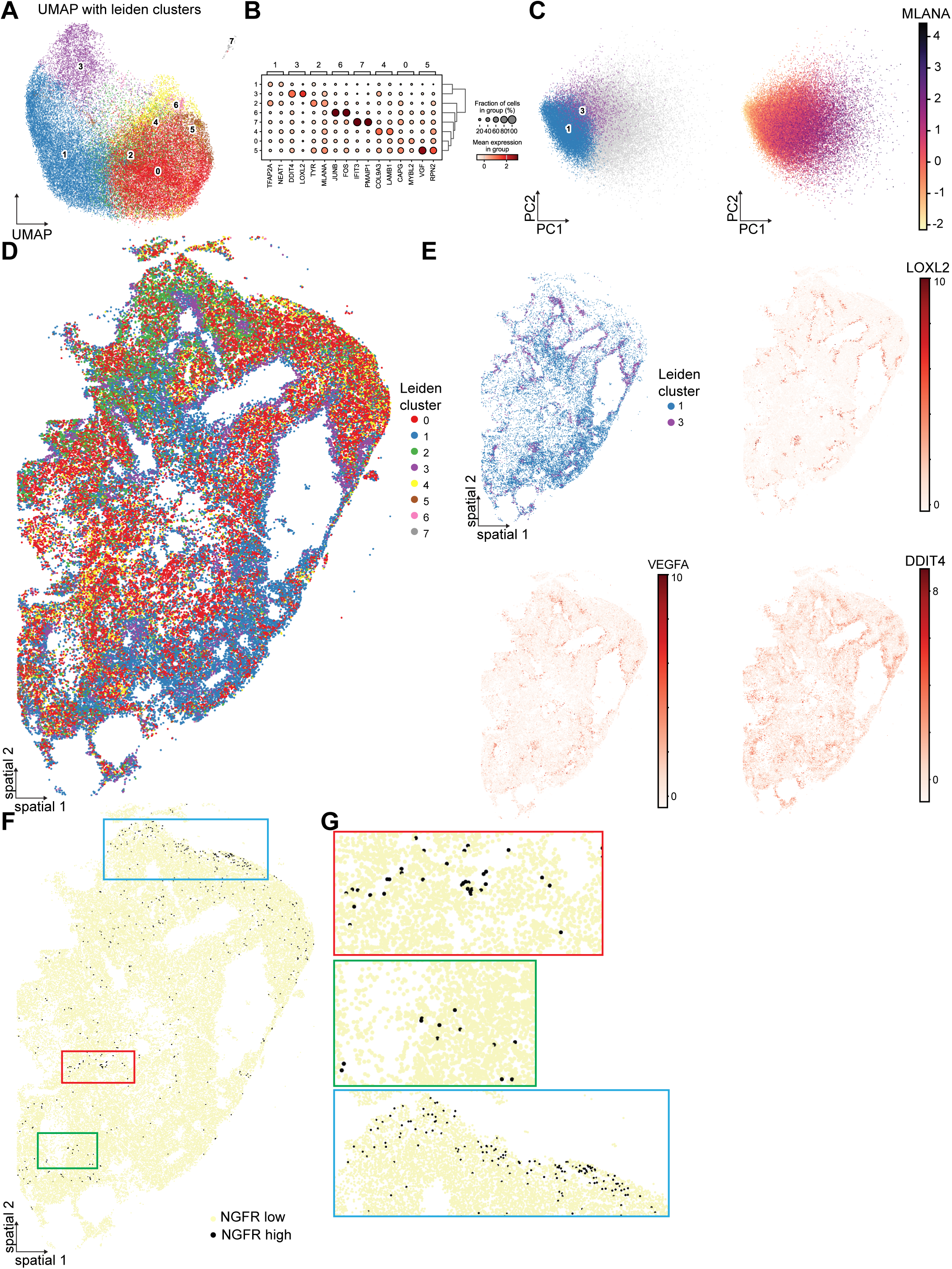
Single-cell spatial transcriptomics of patient-derived xenografts shows intrinsically and extrinsically determined resistant fates. A. Uniform manifold approximation and projection (UMAP) applied to the first 50 principal components for 51897 cells. Cells are colored according to leiden clustering using 10 neighbors and a resolution of 0.6. B. Expression dotplot of the top two marker genes for each cluster across all clusters. Dot size is the fraction of cells in each cluster that express the gene and color is the mean expression of the gene in that cluster. Cluster order on the y-axis is determined by hierarchical clustering. Some leiden clusters have clear, distinct marker genes, such as cluster 3 (DDIT4, LOXL2), cluster 6 (JUNB, FOS), and cluster 7 (IFIT3, PMAIP1). Other clusters, such as cluster 1, have less distinct marker genes. C. Principal component analysis (PCA) of 51897 cells showing the first two PCs. Amount of MLANA expression, a key marker of the melanocytic state, tracks with PC1 (right plot). Clusters 1 and 3 are highly depleted of MLANA and cluster to the left in PC1 (left plot). D. Spatial plot of 51897 cells colored by leiden cluster. Clusters 0, 2, 4, 5, 6, and 7 are dispersed throughout the tissue and are intermixed. Clusters 1 and 3 co-occur around areas of likely necrosis in the tissue section. E. Four spatial plots emphasizing patterns around necrotic regions. Clusters 1 and 3 circumscribe areas of necrosis in the tissue (top left plot). LOXL2, VEGFA, and DDIT4 show expression gradients around the same areas of the tissue (top right, bottom plots). F. Spatial plot showing distribution of NGFR high cells. Cells were binned as either NGFR high or NGFR low by manual thresholding based on the distribution of normalized NGFR values. NGFR high cells are relatively rare and occur in small clusters. NGFR high cells are enriched towards the tissue section edge. G. Zoomed-in regions of the spatial plot F. NGFR high cells occur in small clusters, consistent with transient heritability seen *in vitro* (top, middle). NGFR high cells are not evenly distributed in the tissue and occur more often at the edge of the tissue sample, consistent with some degree of extrinsic fate determination (bottom).

We were curious about the identity and spatial organization of the leiden clusters (clusters 1 and 3) that were specific to the xenograft setting. We first noted that while most cells in the xenograft were high in *MLANA* expression, cells in clusters 1 and 3 were marked by low *MLANA* expression (Figure 2B). Principal component analysis showed that genes corresponding to melanocyte identity, especially *MLANA*, were the drivers of PC1, and that clusters 1 and 3 were low in PC1 (Figure 2C, 2D) Individual marker analysis demonstrated that cluster 1 was marked by *TFAP2A* and *NEAT1*, though notably these genes were not very discriminatory for the cluster (Figure 2B). Instead, its defining characteristic seemed to be its depletion of *MLANA* expression. Cluster 3, in contrast, was strongly marked by *DDIT4* and *LOXL2,* along with low *MLANA* expression (Figure 2B). We next looked at the spatial distribution of leiden clusters throughout the tissue sample (Figure 2D) While most clusters were relatively evenly dispersed throughout the tissue, clusters 1 and 3 formed distinct patterns of zonation (Figure 2D, 2E top left). Clusters 1 and 3 formed relatively large, contiguous areas of expression consisting of dozens to hundreds of individual cells (Figure 2E, top left). These zonation patterns repeatedly formed around regions with very few to no cells, which we took to be necrotic regions of the tissue. We also found spatial patterns of *DDIT4, VEGFA,* and *LOXL2* expression around these necrotic regions (Figure 2E, top right, bottom left and right). Overall, the large numbers of cells involved in this patterning, along with a clear spatial cue, suggest that these resistant fates are extrinsically specified.

In contrast, we found that fates that were found to be intrinsically determined in FateMap were more evenly dispersed throughout the bulk of the tissue. Cluster 7, the interferon cluster, was relatively evenly dispersed and formed small patches of 3-4 cells (Supplementary Figure 7A). Since we know that the interferon fate is intrinsically determined *in vitro*, these small patches of cells presumably came from a single parental cell of origin, though we cannot definitively conclude this because the xenograft lacks lineage barcodes. Cluster 6, which expressed markers of the previously described “stress-like” state, was also evenly dispersed throughout the tissue and occurred in small patches of 3-5 cells (Supplementary Figure 7B). This pattern is suggestive of the stress-like state being similarly intrinsically-determined in the context of a xenograft, though we do not have *in vitro* barcode data to corroborate. In contrast, the distribution of cells with high expression of *NGFR*, which marks the neural crest resistant fate, showed a mixed distribution in the xenograft (Figure 2F). *NGFR*^high^ cells were spread throughout the tissue and occurred in small patches of 3-4 cells, but there also were distinct regions that were enriched for *NGFR*^high^ cells, particularly at the edge of the xenograft (Figure 2F, 2G). The influence of spatial position on the frequency of small patches of *NGFR*^high^ cells suggests that the *NGFR*^high^ fate, though shown to be intrinsically determined *in vitro*, may also be biased depending on the microenvironmental context. Thus, we were able to find both intrinsically and extrinsically determined fates in our single-cell spatial transcriptomic dataset.

## Discussion

In this work, we show that there exists a consensus set of intrinsic resistance programs that associate with the extrinsic tumor microenvironments in melanoma patients treated with targeted therapy. We further show in xenografts that some resistant fates are determined intrinsically and some extrinsically by features of the tumor microenvironment. These are not exclusive possibilities because some fates appear to be determined intrinsically but with a component of bias based on the external environment. Resistant fate specification in this system thus is a complicated mixture of intrinsic and extrinsic determination and may be different for each specific resistance fate.

A common observation in transcriptomics from patient samples is that individual patient signature is the strongest axis of variation and that patient samples tend to cluster separately from each other, potentially due to technical batch effects or biological patient-specific differences. Our work offers a possible reframing of the problem. If one knows the underlying non-genetic programs in the system, these programs can be effectively thought of as a basis set for each transcriptome. This approach avoids the complicated problem of integration and allows one to instead make comparisons across patients by using the relative amount of each non-genetic program. In our case of resistance programs, we used this line of reasoning to calculate the underlying resistance program sets for each sample and only then make comparisons. Of course, this framework requires one to know the basis set *a priori*, which is not always possible.

One major outstanding question is how the underlying genetic background of a cancer cell affects the non-genetic states the cell can access. Our clonal cell line evidence suggests that both driver mutation and broader genetic background impact resistant fate specification, and our patient data shows that certain patients are biased toward certain resistant fates. Atlas-scale single-cell sequencing of resistant tumors from many patients, with or without lineage imputation, could begin to conclusively answer how driver mutation or bulk tumor genome interacts with non-genetic fates. However, since tumors are polyclonal, single-cell genomics combined with transcriptomics would be required to understand how intratumoral genetic heterogeneity drives non-genetic heterogeneity.

Conversely, the role of non-genetic heterogeneity when genetic heterogeneity is available is still unclear. One possibility has to do with timescales. In our system, prior to becoming resistant, a slowly fluctuating non-genetic state drives therapy resistance, but then following acquisition of resistance, the resistant states become “locked in” and are stable over many generations but are not the result of genetic mutations. In patient samples, we see the same resistant states but do not know conclusively whether they are maintained by mutation. One model is that, *in vivo*, the locked-in, non-genetically-encoded states serve as a temporary bridge until the state can be permanently fixed by mutation. It could also be that different mutations allow for different sets of resistant fates to emerge. Such questions could be answered by serial simultaneous profiling of single-cell transcriptomes and genomes.

One limitation of our data is that the direction of causation is impossible to determine. Though we know that *in vitro*, these resistant fates are intrinsically determined, our *in vivo* data lacks lineage information. It may be that differences in the tumor immune microenvironment allow certain resistant fates to emerge and others to die. Or it may be that immune infiltration comes after the resistant fate is determined, with different populations of immune cells attracted to different resistant fates. At least in the context of immunotherapy, non-genetic states can influence T-cell infiltration^32^. In either case, our data clearly demonstrates that resistant fates in targeted therapy interact with the tumor microenvironment.

## Data and Code Availability

All raw and processed data as well as code for analyses in this manuscript can be found at https://www.dropbox.com/scl/fo/3ftnb2a3wf7tkdepukama/AFh1uiJziyjfBBRsX47_jTI?rlkey=vvo8ctnlmjipw018xbc3odj33&dl=0

## Acknowledgments

We thank the Raj lab members for scientific discussion and comments on the manuscript. We thank Ashani Weeraratna and Yash Chhabra for their discussion on melanoma biology. We thank Nancy Zhang for their discussion on statistics. A.R. acknowledges support from a center grant from the Mark Foundation for Cancer Research, NIH Director’s Transformative Research Award R01 GM137425, NIH R01 CA238237, NIH R01 CA232256NIH SPORE P50 CA174523, NIH U01 CA227550, NIH 4DN U01 DK127405. R.H.B. acknowledges support from NIH Training Grant In Computational Genomics T32HG000046 and NIH Medical Scientist Training Program T32GM007170. C.G.T. is supported by the Damon Runyon Cancer Research Foundation, DRG-2465-22. J.A.W. acknowledges support by the NCI Melanoma SPORE (P50CA221703), MD Anderson Melanoma Moonshot, TRANSCEND Cancer Initiative and Platform for Innovative Microbiome and Translational Research (PRIME-TR).

## Author contributions

Conceptualization, A.R.; Methodology, R.H.B, C.G.T., and A.R..; Software, R.H.B., and A.R.; Validation, R.H.B.; Formal Analysis, R.H.B.; Resources, A.R.; Investigation, A.R.; Data Curation, R.H.B., R.L., and A.R.; Writing - Original Draft, R.H.B.; Writing - Review & Editing, A.R., R.H.B., C.G.T., R.L., and J.A.W..; Visualization, R.H.B..; Supervision, A.R.; Project Administration, A.R.; Funding Acquisition, A.R.

## Methods

### Single-cell RNA-seq preprocessing

For input into the cNMF algorithm, we processed each FateMap experiment separately. For each dataset, we started with the cell-by-gene count matrix from CellRanger. When applicable, we applied the same processing done in the original manuscript. Briefly, we required each gene to be present in at least 3 cells, and each cell to have a minimum of 300 features. Other preprocessing parameters were specific to each dataset. All datasets except for the WM989 biological replicate had two technical replicates. Technical replicates were integrated using Seurat’s SCTransform command with the vars.to.regress parameter set to the technical replicate. For the WM989 biological replicate, we applied SCTransform without regressing out a variable. For each sample, we then extracted the ‘counts’ layer for downstream analysis.

### Consensus Non-negative Matrix Factorization

We performed cNMF analysis as previously described^26^. cNMF takes a cell by gene count matrix and produces a matrix of gene expression programs and a matrix with the usage of each program for each cell. The algorithm runs this factorization, which has a degree of randomness, multiple times and performs a meta-analysis of the factorizations to improve robustness and accuracy. Since the user must manually select the number of programs, we ran the factorization independently on each sample over a range of programs from 5 to 13, per the developer’s recommendation. We then selected an optimal number of clusters to maximize the stability and minimize the error of the result while also examining the identity of the gene programs to recover previously identified gene programs in our sample. We finally filtered outlier factorizations and obtained the consensus matrices for each sample.

### Gene expression program overlap

To identify programs that were common across samples, we used a “metaprogram” strategy similar to what has been previously described^28,29^. We took the top 100 marker genes for each NMF gene expression program based on the z score for that program. We then calculated the Jaccard index between the marker genes across all samples, thus finding the overlap between programs. We then clustered the Jaccard index results and considered a Jaccard index of greater than or equal to 0.1 as a significant overlap. Custers that included at least two samples were called metaprograms. We manually inspected the marker genes for the programs that made up each program to assign the metaprogram identity.

### GeoMx data processing

Sequencing results were downloaded and re-processed using slightly different parameters using the manufacturer’s proprietary software package. Default values were used except where otherwise noted. Segments with fewer than 1000 reads, less than 80% aligned reads, or less than 50% sequencing saturation were excluded from downstream analysis. Probes were excluded if the ratio of the geometric mean of the probe in all segments to the geometric mean of the probe in the target was less than or equal to 0.1 or if the probe failed the Grubbs outlier test in 20% or more of samples. Finally, we kept targets that exceeded a threshold (higher of limit of quantification or count of two) in at least 2% of samples. We exported the filtered count data for downstream analysis. Finally, we had a pathologist manually annotate the hematoxylin and eosin eosin-stained and fluorescent-stained regions for a number of features reported in supplementary table 2.

### Non-negative least squares metaprogram deconvolution of GeoMx data

To perform non-negative least squares (nnls) deconvolution, we needed to construct a signature matrix and a target matrix in the same units. For the signature matrix, we imported all the genes that were markers for the metaprograms identified above. We then imported the gene usage matrices for each experiment in units of CPM. We filtered these usage matrices to include only the marker genes from the metaprogram. We constructed the signature matrix by taking the mean gene usage across individual programs from each FateMap experiment for each metaprogram. Finally, we filtered genes from the signature matrix that appeared in the safeTME signature matrix in order to minimize potential confounding from genes expressed in the tumor microenvironment. We constructed the target matrix by doing upper quartile normalization and CPM calculation of the filtered GeoMx count data using edgeR. We then performed nnls deconvolution of each GeoMx sample on the common genes to the target and signature matrix using the ‘nnls’ command from the nnls package.

### Likelihood and empiric p-value calculation for non-negative least squares metaprogram deconvolution

To assess the quality of the deconvolution using metaprograms, we compared the likelihood of the deconvolution to a shuffled null distribution of log-likelihoods under a negative binomial distribution. We first found the gene expression values that would be predicted from our deconvolution by multiplying the signature matrix by the transverse of the deconvolution matrix. This predicted gene expression matrix contained values of 0, which are not possible under a negative binomial and would give an indeterminate log-likelihood value. To account for this, we added an adjustment value to each predicted gene expression value that was equal to the smallest non-zero predicted gene expression value in the matrix. We then calculated the log-likelihood of observing the true gene expression value, rounded to the nearest whole number, under a negative binomial distribution with mu equal to the adjusted predicted gene expression value from the deconvolution. Since we had no estimate for the size parameter of the negative binomial, we calculated the log-likelihood with size parameters of 5, 10, and 20. This produced a log-likelihood for each gene in each sample. We finally summed the log-likelihood of all genes in a sample to get the total log-likelihood for each sample.

To calculate an empiric p-value for the deconvolution, we constructed a null distribution using a shuffling approach. We randomly shuffled the gene labels of the signature matrix, deconvoluted the data, and calculated the log-likelihood of observing the gene expression values as described above. We repeated this shuffling procedure 1000 times and thus found a null distribution of log-likelihoods for each sample. We calculated a one-tail empiric p-value for each sample by finding the number of null log-likelihoods that were greater than the true log-likelihoods. We report these p-values in supplementary table 1.

### Tumor microenvironment SpatialDecon of GeoMx data

We performed tumor microenvironment cell deconvolution of the GeoMx data using the manufacturer’s software package SpatialDecon^33^. The package contains a tumor microenvironment deconvolution signature matrix, safeTME, that is specifically designed to exclude genes that are commonly expressed in tumor cells. We further filtered this signature matrix to exclude genes that were called as variable genes in therapy-resistant WM989 and WM983b cells. Performed the deconvolution using the manufacturer’s recommended settings except for as noted below. We estimated the background from negative targeting probes using the ‘derive_GeoMx_background’ command. We then used the estimated background to perform the deconvolution using the ‘spatialdecon’ command on the upper-quantile normalized data using our filtered safeTME signature matrix and nuclei counts from the machine with additional parameters ‘align_genes = TRUE’, ‘cellmerges = safeTME.matches’, ‘n_tumor_clusters = 5’, and ‘is_pure_tumor = grepl(“S100”, colnames(dsp@assayData$exprs))’. We exported the deconvolution for visualization and further analysis.

### Comparison of tumor microenvironment and metaprogram deconvolutions

Across all regions of interest, we performed pairwise correlation analysis of each resistance metaprogram against each immune cell type using the deconvolutions described above. We did so by calculating a Spearman correlation and associated p-value for each pair using the ‘cor’ and ‘cor.test’ functions, respectively, with parameter method = ‘spearman.’ We selected a p-value cutoff of 0.01 to denote statistically significant associations.

### Spatial transcriptomics with Spatial Genomics platform

#### Panel design

We designed a custom panel based on the SeqFISH technology from SpatialGenomics. The panel was chosen based on expert curation of the literature and expression in the *in vitro* FateMap datasets. The final panel contained 427 genes, of which 399 were detected using barcoded seqFISH imaging and 28 were identified sequentially via single-molecule FISH. The panel was designed in collaboration with SpatialGenomics and synthesized by them. The panel can be found in supplementary table 3.

#### Sample preparation and imaging

We used patient-derived xenograft (PDX) samples that had been collected in a previous study^34^. We prepared and ran a PDX sample using the manufacturer’s protocol from SpatialGenomics. We cryosectioned a 5µm section onto a proprietary Spatial Genomics slide. We then followed the frozen tissue sample preparation method. We fixed the sample using 4% formaldehyde in PBS and then dehydrated and permeabilized with 70% ethanol. We cleared the sample with the proprietary clearing solution for 5 minutes, rinsed in 70% ethanol, and assembled the flow cell. We then washed the sample in the primary wash buffer, denatured primary probes at 90C for 3 minutes, and hybridized the primary probes to the sample at 37C overnight. The next day, we washed the sample with a primary wash buffer, stained the nuclei with a DAPI-containing staining solution, and added a rinse buffer to the sample. We then loaded the sample into the machine for rounds of secondary hybridization and readout with our custom-designed expression kit.

#### Image processing and upstream analysis

All image processing was done either on the SpatialGenomics machine or with the SpatialGenomics proprietary software. Raw images were aligned across multiple hybridizations to form composite images. Spots were detected by manually thresholding each channel to maximize spot detection while minimizing background detection. When manually thresholding, we found that a number of genes detected by smFISH were either poorly detected or had high background such that spots could not be called. We noted these nine genes and removed them from the analysis. The transcript identities were then decoded, and nuclei were segmented using a machine-learning algorithm on the DAPI image. A mask was formed from the segmented nuclei that included the nucleus plus a dilation around the nucleus. Individual transcripts were then assigned to individual cells, thus yielding a cell-by-gene count matrix that was exported to Python for downstream analysis.

### Spatial transcriptomics unsupervised clustering and spatial analysis

Cell-by-gene count matrices and nuclei xy-locations were imported into Python and analyzed using the Squipy package (see the accompanying .yml file for the full conda environment)^35^. One challenge was filtering out neighboring mouse cells from our analysis. To do so, we implemented multiple filtering steps prior to unsupervised analysis and clustering of cells. First, we filtered cells based on the nuclear area, only keeping those that were between 2000 and 60000 pixels in area. These thresholds were chosen both by manual inspection of the images to compare the relative size of human and mouse cells and by looking at the distribution of cell areas. We next excluded cells that contained 50 or fewer transcripts, that contained fewer than 21 unique genes, and that contained fewer than 20 unique genes detected by barcoding rather than smFISH. Our reasoning was that mouse cells would have fewer overall transcripts and, in particular, have fewer barcoded transcripts because genes detected by the barcode method require multiple spots to be detected over multiple rounds of imaging.

We then normalized cell counts by the area of the nucleus, as has been recently recommended for this type of image-based spatial transcriptomic data^36^. We then scaled the data and performed principal component analysis (PCA). We then found neighbors using the ‘pp.neighbors’ command with parameters n_neighbors = 10 and n_pcs = 50. We then performed Uniform Manifold Approximation and Projection (UMAP) analysis using the ‘tl.umap’ command with parameter min_dist = 0.01. Finally, we performed leiden clustering using the ‘tl.leiden’ command with parameter resolution = 0.6. We found marker genes for the leiden clusters using the ‘tl.rank_genes_groups’ using both the logistic regression and Wilcoxon strategy. We thresholded cells as being ‘NGFR_high’ if the expression value was greater than or equal to 3, based on manual inspection of the expression distribution. Since some of these processing steps rely on random seeds, we exported these processing steps to a .h5ad file before any plots were made to maximize reproducibility. All plots were made using built-in Scanpy and Squidpy plotting functions.

## Supplementary figure captions

**Supplementary figure 1.**
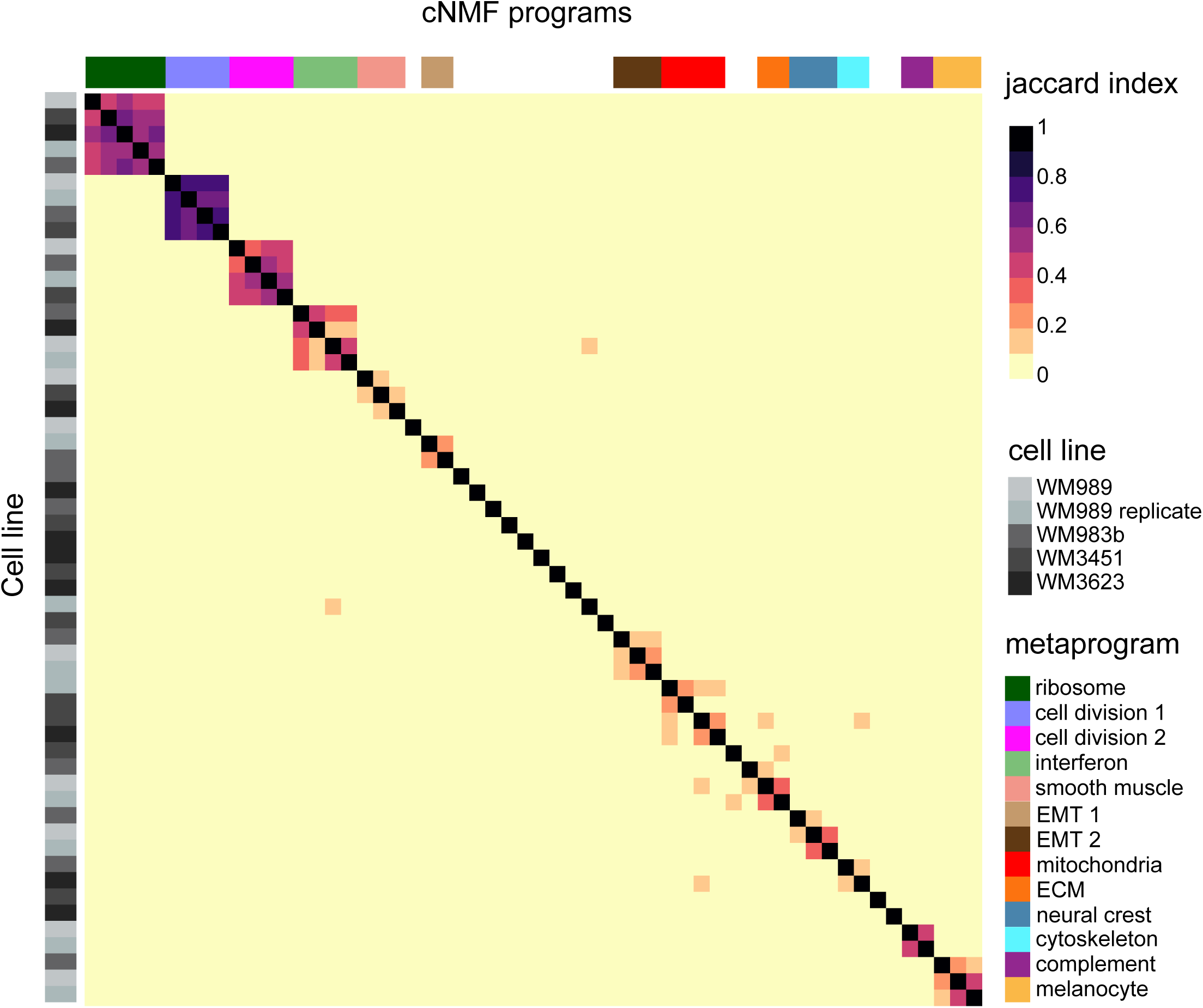
Heatmap of Jaccard index of top 100 marker genes for each cNMF program across all samples. cNMF programs were identified separately for each sample. We extracted the top 100 marker genes for each program and calculated the Jaccard index between each pairwise combination of gene programs. We then performed hierarchical clustering on the results and plotted the resulting heatmap of Jaccard indices. We defined a metaprogram as a gene program that had at least a 0.1 Jaccard index between two samples. We named each metaprogram according to expert curation of the genes that appeared in all samples included in that metaprogram.

**Supplementary figure 2.**
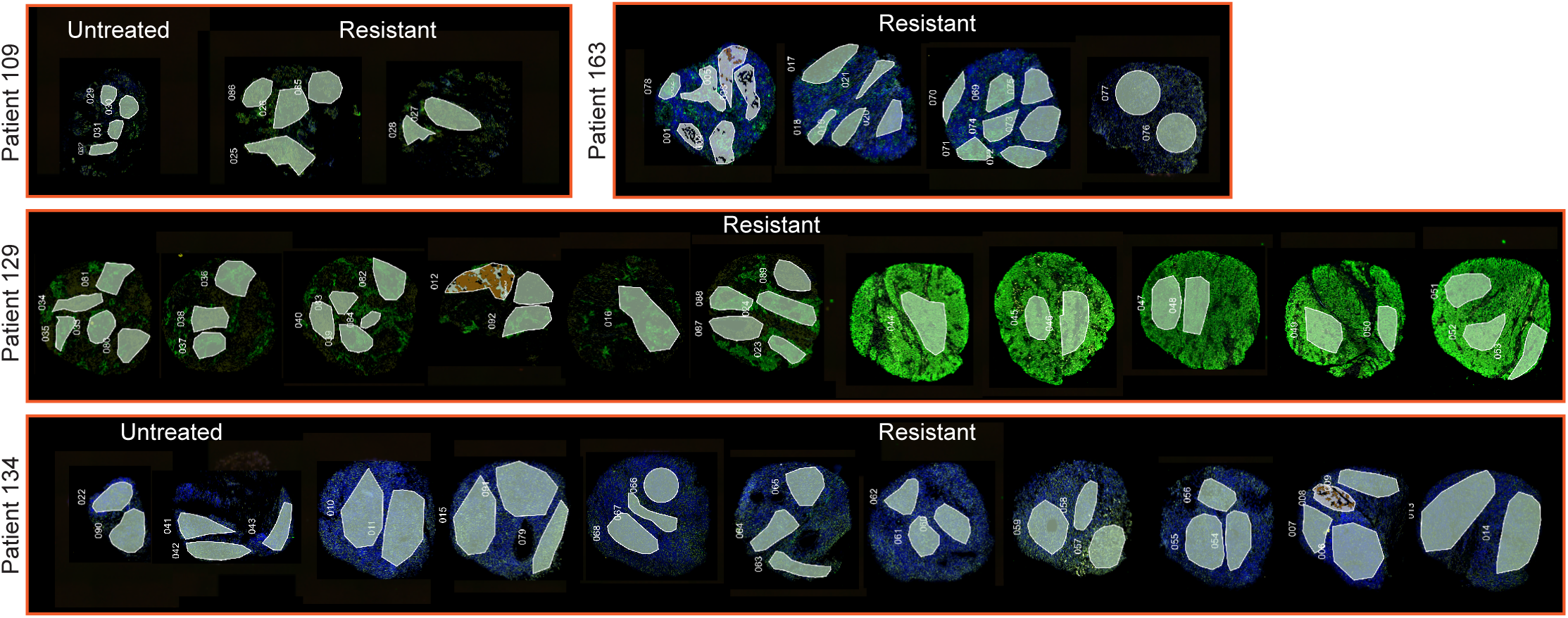
All patient sample plugs with annotated ROIs. All 29 punch biopsies from four patients, two with matched pre-treatment samples, that were sequenced using the GeoMx Digital Spatial Profiler platform for spatial transcriptomics. 93 total ROIs were selected for sequencing based on staining for DNA (SYTO 13, blue), S100B (green), and CD45 (red).

**Supplementary figure 3.**
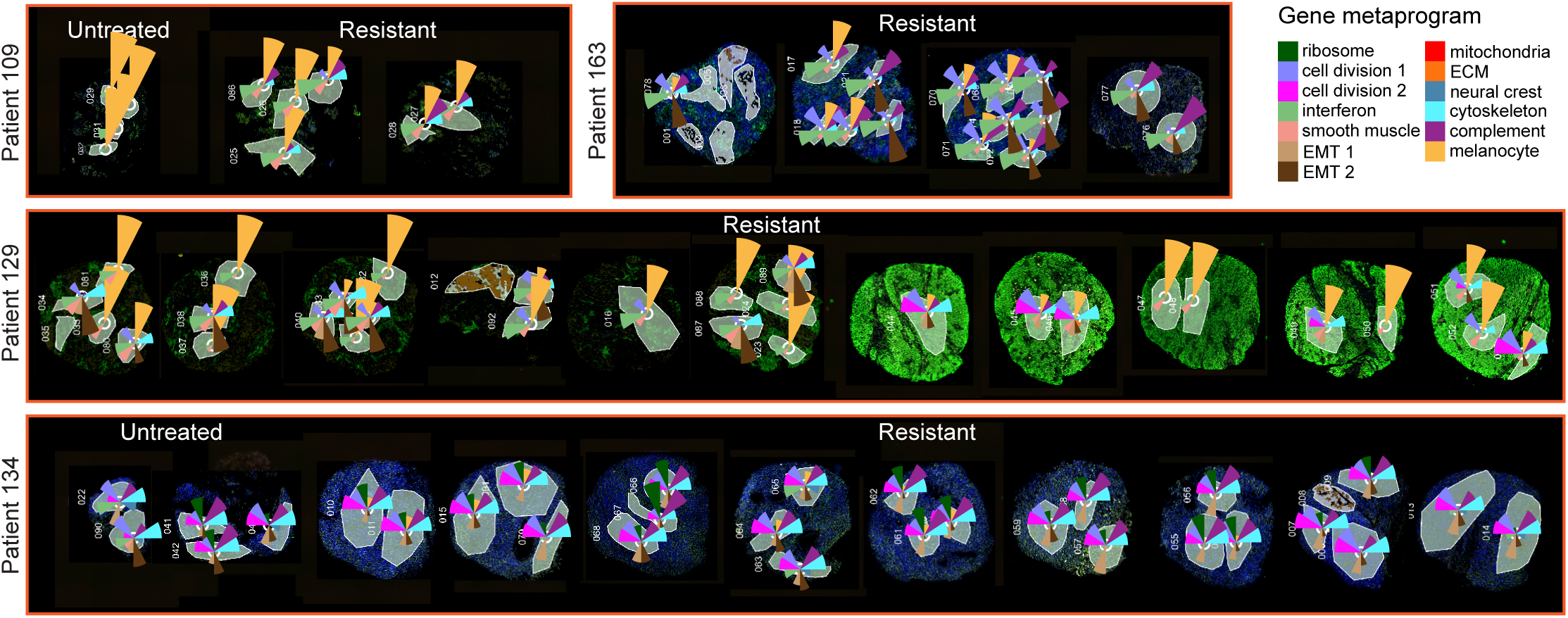
Non-negative least squares deconvolution of all full GeoMx ROIs using the twelve metaprograms identified from *in vitro* cNMF analysis. For each full ROI (that was not split based on fluorescence signal and separately sequenced), we performed non-negative least squares (nnls) deconvolution on the whole transcriptome using the expression values of twelve listed metaprograms as the signature matrix. For each ROI, we normalize the metaprogram deconvolution to one and then plot the relative amount of each metaprogram as the area of the pie wedge. The distribution of metaprogram expression shows multiple metaprograms emerge within a single patient, although each patient may have a bias towards a particular distribution of these programs.

**Supplementary figure 4.**
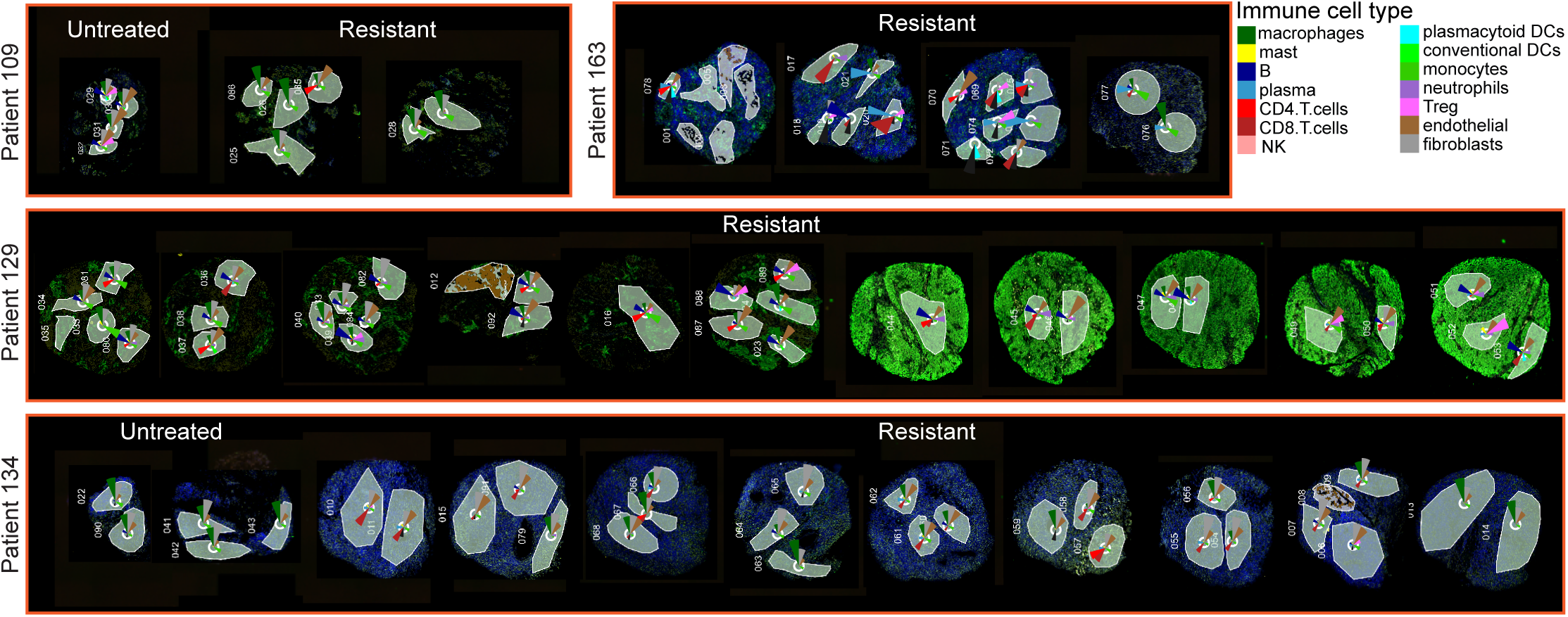
Spatial deconvolution of all full GeoMx ROIs using the manufacturer’s immune cell signature matrix. For each full ROI (that was not split based on fluorescence signal and separately sequenced), we performed deconvolution on the whole transcriptome using command ‘spatialdecon’ from the SpatialDecon package and using the ‘safeTME’ immune cell signature matrix from the that we additionally filtered to remove genes that were variable genes in the WM989 and WM983b datasets to minimize the influence of transcripts from tumor cells in the estimation of immune cell amounts. For each ROI, we normalize the immune cell deconvolution to one and then plot the relative amount of each immune cell type as the area of the pie wedge.

**Supplementary figure 5.**
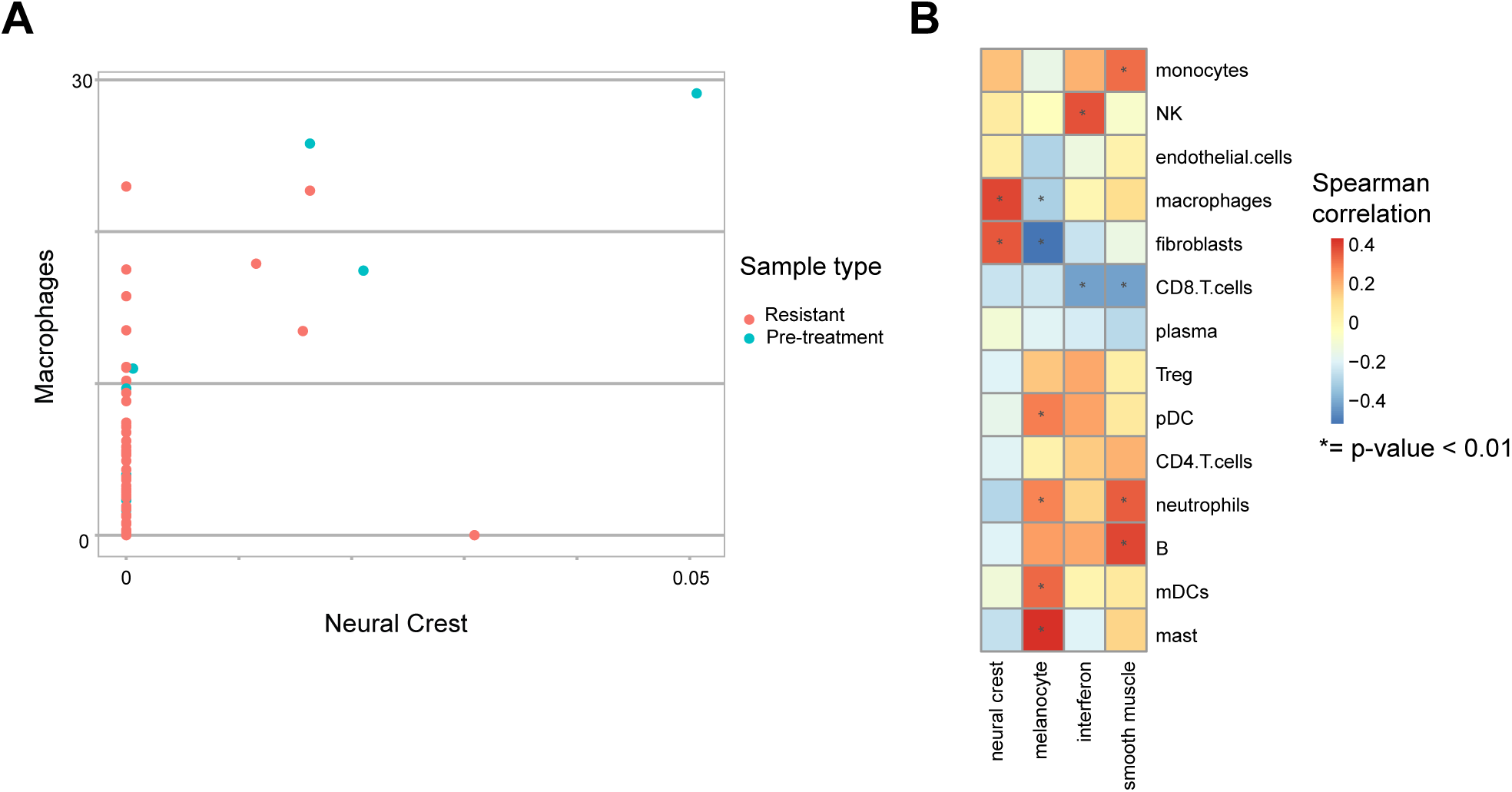
Association of resistance program and immune signature including pre-treatment ROIs. A. A small but statistically significant association between the neural crest resistance program and macrophages emerges only when including pre-treatment ROIs. Each point is an individual ROI, the x-axis is the ROI’s deconvolution for the neural crest resistance fate, and the y-axis is the deconvolution for the macrophage signature. Color is whether the ROI came from a pre-treatment or resistant punch biopsy. B. Heatmap of Spearman correlation values for resistance program compared to immune cell signature for tumor-containing ROIs. The associations change slightly when including pre-treatment samples in the analysis including a new positive correlation between the neural crest program score and the fibroblast and macrophage signature.

**Supplementary figure 6.**
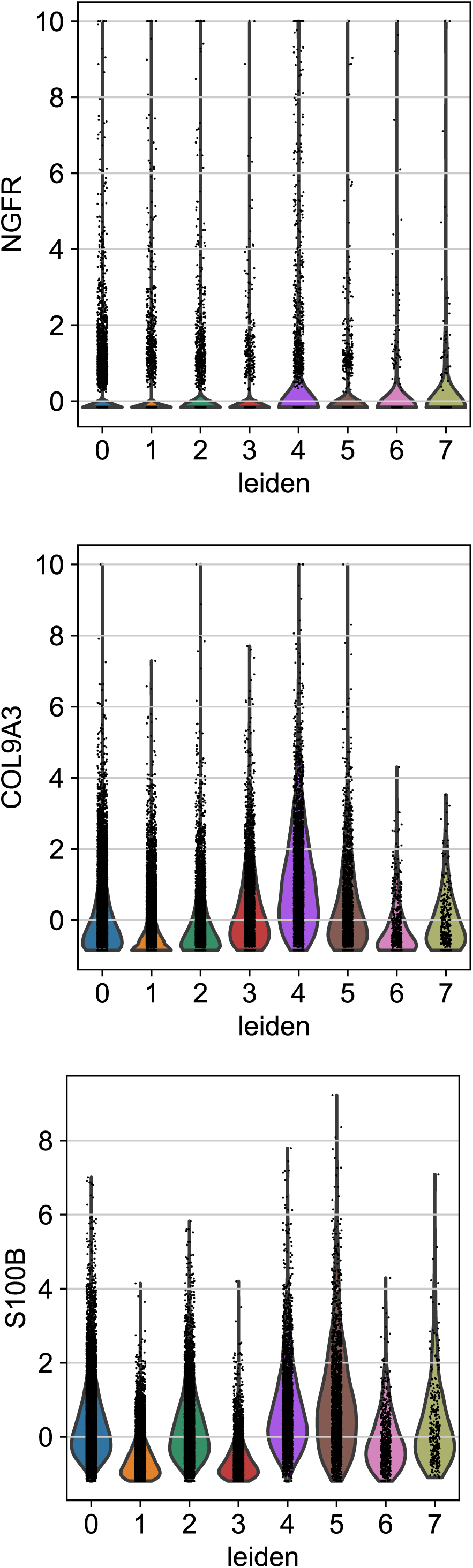
Neural crest resistant fate marker expression across leiden clusters. Violin plots of the expression of markers of the neural crest resistant fate across unsupervised leiden clusters. *In vitro*, *NGFR*, *COL9A3*, and *S100B* are sensitive and specific markers of the neural crest resistant fate and their expression is relatively confined to a single unsupervised cluster. These three genes are enriched in cluster 4 of the PDX sample. However, unlike *in vitro*, all three genes are expressed in multiple leiden clusters, and *COL9A3* and *S100B* but not *NGFR* score as top markers of cluster 4.

**Supplementary figure 7.**
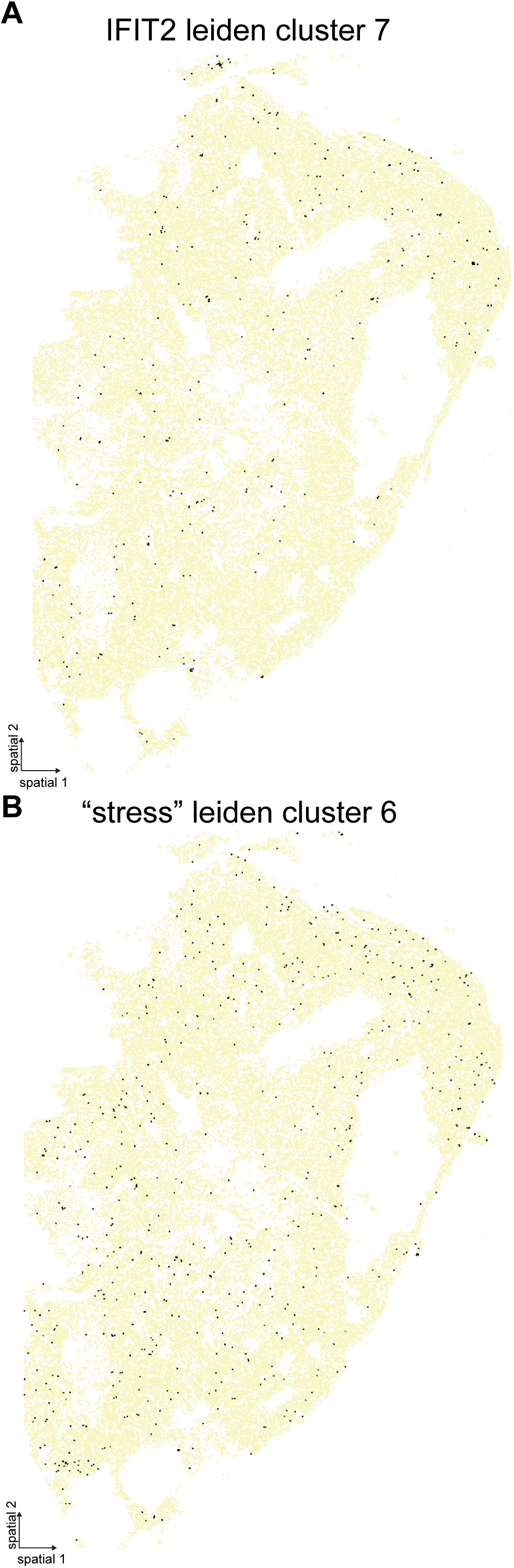
Spatial distribution of stress-like and IFIT2 clusters. A. Spatial plot with cells in cluster 6, the “stress” cluster, as black dots and cells in all other clusters as pale dots. Cluster 6 is relatively evenly dispersed throughout the tissue but also occurs in small clusters of 3-5 cells suggesting a common parental cell of origin. B. Spatial plot with cells in cluster 7, the IFIT2 positive cluster, as black dots and cells in all other clusters as pale dots. Like cluster 6, cluster 7 is also evenly dispersed throughout the tissue but notably occurs in clusters of 3-5 cells, again suggesting a common parental cell of origin.

